# Temporal changes in the nasal microbiota and host antimicrobial responses to intranasal mupirocin decolonisation: Observations in healthy staphylococcal carriers

**DOI:** 10.1101/126375

**Authors:** Su-Hsun Liu, Yi-Ching Tang, Yi-Hsiung Lin, Kuan-Fu Chen, Chih-Jung Chen, Yhu-Chering Huang, Leslie Y. Chen

## Abstract

**Background:** How intranasal mupirocin decolonisation affects the human nasal microbiota remains unknown. To characterize the temporal dynamics of the nasal microbial community in healthy staphylococcal carriers in response to intranasal mupirocin decolonisation, we serially sampled the anterior nares of four healthy carriers to determine the nasal microbial profile via sequencing of bacterial 16S ribosomal DNA.

**Results:** Before decolonisation, the nasal microbiota differed by the initial, culture-based staphylococcal carriage status, with *Firmicutes* (54.1%) and *Proteobacteria* (75.8%) dominating the microbial community in the carriers and the noncarrier, separately. The nasal microbiota lost its diversity immediately after decolonisation (Shannon diversity: 1.33, 95% confidence interval [CI]: 1.06-1.54) as compared to before decolonisation (1.78, 95%CI: 0.58-1.93). The initial staphylococcal carriage status, expression levels of human neutrophil peptide 1, and sampling times were major contributors to the between-community dissimilarities (*P* for marginal permutation test: .014) though of borderline significance when considering data correlation (*P* for blocked permutation test: .047) in both nonmetric multidimensional scaling and constrained correspondence analysis. Results of univariable and multivariable differential abundance analysis further showed that, in addition to Staphylococci, multiple genera of Actinobacteria and Proteobacteria were differentially enriched or depleted by mupirocin use.

**Conclusions:** Mupirocin could affect both Gram-positive and Gram-negative commensals along with altered host antimicrobial responses. How the nasal microbiome recovered after short-term antibiotic perturbation depended on the initial staphylococcal carriage status. The potential risks associated with loss of colonisation resistance need to be considered in high-risk populations receiving targeted decolonisation.

## BACKGROUND

In the U.S., 80% of *S. aureus* infections occurred in the healthcare setting. Among surgical patients and patients newly-admitted to the intensive care unit,[1] nasal staphylococcal carriage has been consistently shown to increase infection risks by 2-12 folds at surgical sites (SSI), in the bloodstream, or at catheter exit sites.[2] While targeted decolonisation with intranasal mupirocin calcium ointment, either alone or in a bundle, can effectively reduce *S. aureus*-associated SSIs among certain carriers,[3] the potential increase in bacterial resistance upon prolonged or widespread implementation of antibiotic decolonisation cannot be over-estimated.[4]

In a recently published meta-analysis, universal and targeted decolonisation appeared to be similarly effective on reducing *S. aureus* infections in non-surgical healthcare settings.[5] However, the authors noted that 5 out of 14 studies reported the emergence of mupirocin resistance.[5] Grothe and colleagues also cautioned the development of drug resistance in specific patients populations for whom repeated or prolong decolonisation may be clinically indicated.[6] Cumulative evidence from animal models and clinical studies have suggested that, the human microbiota constitutes an essential microenvironment that may promote or inhibit colonisation of certain bacterial species by inter-species or host-pathogen interactions.[7] Longitudinal observations of infants exposed to antibiotics revealed that the resulting dysbiosis of the gut microbiota could be long-lasting and might predispose to the later development of allergic or atopic phenotypes, obesity, and other autoimmune disease.[8] In adults, repeated exposure to antibiotics would eventually lead to a stable yet different-from-baseline microbiome in the distal gut; the functional consequences remained to be determined.[9]

While staphylococcal carriers are known to harbour a characteristically distinctive microbial profile in the anterior nares as compared to noncarriers,[10, 11] there is a paucity of literature on how antibiotics may affect the human nasal microbiota immediately and in a short-term fashion as in the post-decolonisation surgical or ICU setting. Understanding the impacts of antibiotics on the nasal microbiome and how the host immunity reacts, can support evidence-based decolonisation policies.[4] Therefore, we sought to characterize temporal changes in the nasal microbial community by following healthy staphylococcal carriers up to three months after intranasal mupirocin decolonisation. We also attempted to quantify antimicrobial responses of the host skin associated with a perturbed nasal microbiota.

## METHODS

### Study design and data collection

The current study extended from a parent cohort study on healthy staphylococcal carriers, the study protocol of which was previously described.[12] Briefly, we sampled the anterior nares of female carriers twice per week for six consecutive menstrual cycles. Four of the 15 participants requested for a standard course of topical decolonisation at the end of the study, at which time (April 2015) there were no clinical consensus on screening or decolonising staphylococcal carriage in Taiwan.

After obtaining written informed consent, we sampled anterior nares of each participant before (T0) and after a five-day course of mupirocin at 3, 17, 31, 60 and 90 days (T1-T5). The decolonisation strategy was in accordance to the clinical guidelines issued by the Infectious Diseases Society of America.[13] At each sampling visit, the research assistant used one Copan^®^ culture swab for culture-based identification of *S. aureus*; the other sterilized cotton swab for microbial 16S ribosomal RNA gene sequencing; still another sterile cotton swab for gene expression quantification. The sampling order of the three swabs remained the same for all subjects throughout the observation period. The Institutional Review Board at Chang Gung Medical Foundation reviewed and approved the addendum study protocol and the consent form.

### Laboratory procedures

#### Microbiology study

According to the current guideline, we performed microbial isolation within 48 hours of swab collection; MRSA isolates were determined by results of the disk diffusion method that demonstrated resistance to both Penicillin and Cefoxitin.[14, 15]

#### RNA extraction and transcript quantification

We stored nasal swabs at -80°C before using TRIzol Reagent^®^ to extract total RNA. To quantify expression levels of selected host antimicrobial peptides (AMP) including HNP1 (*DEFA1*),[16] RNase 7 (*RNASE7*),[17] and HBD3 (*DEFB103A*),[18] we used the reversely-transcribed cDNA as a template for RT-qPCR using Roche Light Cycler 480^®^. In accordance with the literature, we chose glyceraldehyde-3-phosphate dehydrogenase (GAPDH) gene as the reference gene,[19] and median log_2_-fold changes (with inter-quartile range [IQR]) for statistical comparisons using median regression models with robust variance estimation in Stata (version 13) to account for repeated measures.[20]

#### Targeted amplicon preparation and sequencing

We used the primer pair 27F-534R for targeted amplification and pair-end sequencing of the variable regions V1-V3 of 16S ribosomal cDNA [21] using Illumina MiSeq^®^ (2×300 pair-end reads). Due to the poor quality of Read 2 sequences, the following analysis used only Read 1 sequences. There were totally 1,958,527 sequences obtained, with an average of 81,605 sequence reads per sample. We employed the ‘closed_reference’ approach for taxonomy assignment against the Greengenes database (Version 13.8) [22] via the UCLUST algorithm [23] at a threshold of 97% similarity. After quality checking procedures according to the QIIME pipeline (version 1.9.1),[24] we performed additional data management and subsequent statistical analysis in R (Version 3.2.4). [25]

### Statistical analysis

#### Community richness, evenness and diversity

We compared changes in the microbial community structure using the following ecological metrics: *alpha-diversity estimates*, including the observed number of operational taxonomy units (OTU) and the Chao1 index; *beta-diversity estimates*, including Shannon diversity (D_Shannon_) and Simpson diversity (D_Simpson_) indices for quantifying community richness and diversity simultaneously.[26] We also calculated Shannon evenness (E_Shannon_) and Simpson evenness (E_Simpson_) measures to quantify richness-independent evenness of component members.[27] We applied Mann-Whitney U test, Wilcoxon signed rank test or median regression models with robust variance estimation whenever appropriate. In sensitivity analysis, we further computed the above-mentioned indices on rarefied data using coverage (sequence reads)-based rarefaction at a subsampling depth of 41,698 reads (without replacement).[26, 27]

#### Community similarity and dissimilarity

We identified major OTUs (≥ 5% in each sample) concurrently shared by all participants and applied the Morisita-Horn similarity (overlap) index (SI) to quantify pairwise, OTU-by-OTU shared information among all participants or by the initial carriage status.[27] We further performed nonmetric multidimensional scaling (NMDS) to determine “environmental” variables contributing to the between-sample dissimilarities by use of the Morisita-Horn dissimilarity index.[27] Covariates considered included sample-level factors (different sampling times, pre- versus post-mupirocin, days since the start or completion of mupirocin use); subject characteristics (baseline carriage status); relative expression levels of AMPs at each visit. We then constrained the analysis on community dissimilarities that could be best explained by selected host characteristics using constrained correspondence analysis (CCA).[26] Lastly, we performed differential abundance analysis to identify specific OTUs that were differentially enriched or depleted by the baseline staphylococcal carriage status or by expression levels of AMPs. We chose Wald test to compare geometric mean-normalized count data and specified the significance level of Benjamini-Hochberg adjusted P-values at 0.001.[28, 29] We considered a two-tailed, statistical significance at 0.05 unless otherwise specified; p-values were derived from 1000 permutations.

## RESULTS

The included healthcare workers aged between 22 and 42 years, and have spent at least one year at their current job position (Supplementary Table S1). In total, we collected 24 nasal swabs, with six from each participant. Before mupirocin decolonisation (T0), three women carried *S. aureus* (75%) based on culture results. Despite transient detections of *S. aureus* after decolonisation at 3, 5 and 9 weeks, the clearance rate of staphylococcal carriage at post-decolonisation 90 days (or 13 weeks, T5) was 100%.

According to results of relative quantities of selected AMPs, expression levels of RNase7 were undetectable in multiple samples and thus not shown. While there were no differences in HBD3 levels by staphylococcal carriage (*P*: .439, Table 1), there was a 1.59-fold (95% CI: 1.08-2.36) change in HBD3 expression over time (*P* for trend: .022). By contrast, levels of HNP1 expression showed no obvious temporal pattern but were generally lower in carriers than in the noncarrier, either without (*P*: .033) or with (*P*: .016) adjustment for sampling times (Table 1).

**Table 1.** Fold changes in quantities of transcripts from selected antimicrobial peptide genes relative to the baseline (pre-treatment) by the initial carriage status

| AMP | All (n=24) |  | Non-carrier<br>(n=6) | Carrier (n=18) |  |
| --- | --- | --- | --- | --- | --- |
|  | Median | (IQR) |  | Median | (IQR) |
| hBD3 |  |  |  |  |  |
| 1 wk | 2.91 | (0.59, 7.52) | 4.76 | 1.07 | (0.11, 10.3) |
| 3 wk | 2.30 | (1.83, 2.55) | 1.52 | 2.48 | (2.13, 2.61) |
| 5 wk | 10.7 | (5.35, 23.9) | 3.65 | 14.3 | (7.06, 33.6) |
| 9 wk | 4.14 | (1.23, 33.4) | 0.95 | 6.77 | (1.52, 59.9) |
| 13 wk<sup>a</sup> | 6.17 | (1.75, 12.9) | 1.41 | 10.3 | (2.08, 15.5)<sup>b</sup> |
| HNP1 |  |  |  |  |  |
| 1 wk | 0.67 | (0.13, 1.36) | 1.61 | 0.22 | (0.04, 1.11) |
| 3 wk | 0.36 | (0.22, 0.84) | 1.27 | 0.32 | (0.12, 2.22) |
| 5 wk | 0.64 | (0.17, 2.48) | 4.00 | 0.32 | (0.02, 3.33) |
| 9 wk | 0.20 | (0.09, 1.95) | 3.68 | 0.17 | (0.01, 4.44) |
| 13 wk<sup>c</sup> | 0.87 | (0.41, 2.00) | 3.02 | 0.76 | (0.06, 5.55)<sup>d</sup> |

Abbreviations: AMP, antimicrobial peptide; IQR, interquartiel range; hBD3, human beta-defensin3; HNP1, human neutrophil peptide 1; wk, week
a. P for trend: 0.022
b. No statistical difference by carriage status, with (P: 0.497) or without (P: 0.439) adjusting for study visit
c. P for trend: 0.141
d. Statistical difference by carriage status, with (P: 0.016) or without (P: 0.033) adjustment for study visit.

### Community structure of the nasal microbiome

Details of sample sequencing metrics are shown in Supplementary Table S2. Before decolonisation, the dominant phylum was *Proteobacteria*, mostly of unclassified *Oxalobacteraceae* in the only noncarrier (73.0%, Supplementary Figure S1) whereas Firmicutes were predominant colonizing microbes in carriers (T0, Figure 1). Immediately following decolonisation (<1 week, T1), Gram-negative *Oxalobacteraceae* took dominance in all subjects with a relative abundance ranging from 49.8% to 93.7% before *Actinobacteria* (mainly *Corynebacterium*) and *Firmicutes* (unclassified *Bacillaceae* and *Staphylococcus*) gradually re-emerged in the noncarrier and carriers, respectively (Figure 1).

**Figure 1.**
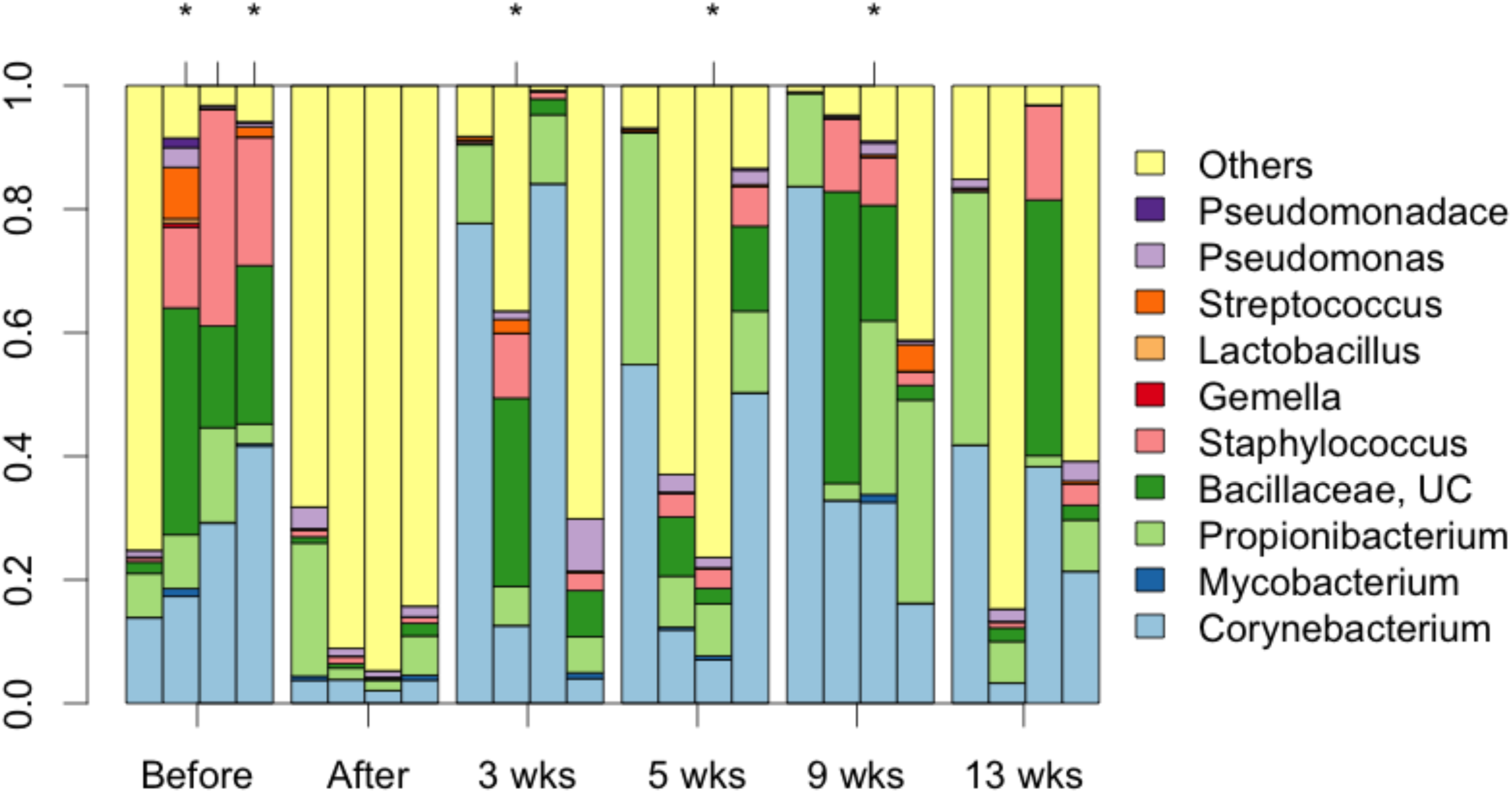
Relative abundance of most abundant 10 genera at each sampling time.

### Community richness, evenness, and diversity

Overall, we identified 341 unique OTUs in 24 swab samples, with an estimated Chao1 index of 361.6 (standard error [SE]: 10.5, Table 2). The degree of community richness was generally greater before decolonisation whereas the community diversity showed the opposite trend (Supplementary Figure S2). The within-sample diversity as quantified by D_Shannon_ (median: 1.69, IQR: 1.20-1.87) and D_Simpson_ (median: 0.66, IQR: 0.46-0.76) represented 50.2% and 56.1% of the overall diversity (2.39 and 0.82), and was higher after (T1-T5) than before mupirocin decolonisation (T0) though the difference was not statistically significant after correcting for within-group correlation (Table 2).

**Table 2.**
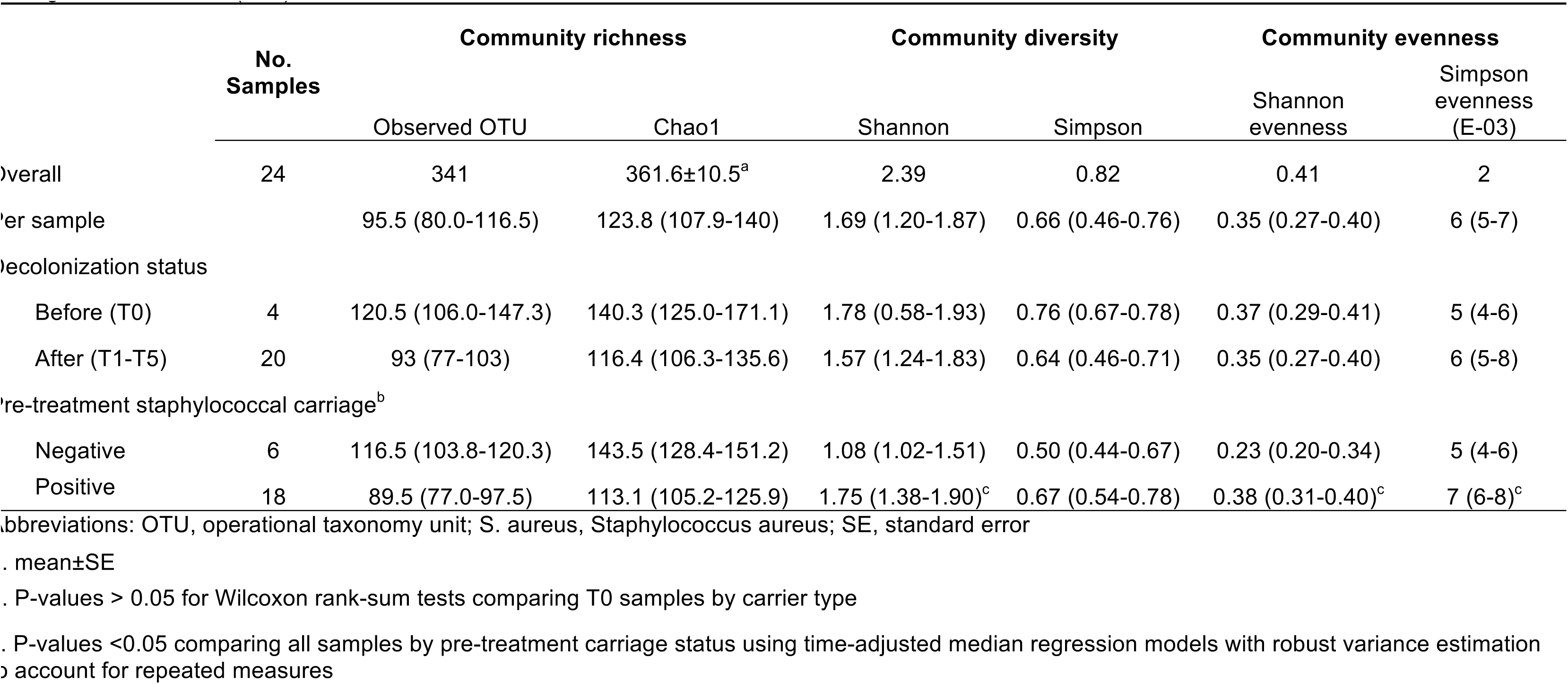
Diversity estimates for the community composition of the nasal microbiota: overall, by sampling time, and by pre-treatment staphylococcal carriage status, median (IQR)

Despite that carriers demonstrated a comparable community richness to that in the noncarrier, an increased evenness (E_Shannon_: 0.38 vs. 0.23, *P*<.001) and diversity measures (D_Shannon_: 1.75 vs. 1.08, *P*: .003) appeared to be associated with the initial staphylococcal carriage status (Table 2). When using rarefied data, richness estimates ranked the community samples comparably and diversity estimates also followed a similar temporal pattern (data not shown).

### Community similarity and compositional differences over time

The proportion of shared OTUs before decolonisation (T0) (20.1%) remain relatively unchanged over time (*P* for trend: .411) but the overall Morisita-Horn overlap estimate increased over 50% immediately after mupirocin use (0.927, 95%CI: 0.925-0.929) before returning to the baseline level with a minimal decrease of 8% at 13 weeks post-mupirocin use (Table 3). Results of between-group comparisons of shared OTUs (%) revealed a resemblance between noncarrier and carriers samples within 5-6 weeks of mupirocin use as compared to that at baseline (T0) or after 8 weeks (T4 & T5, Table 3).

**Table 3.**
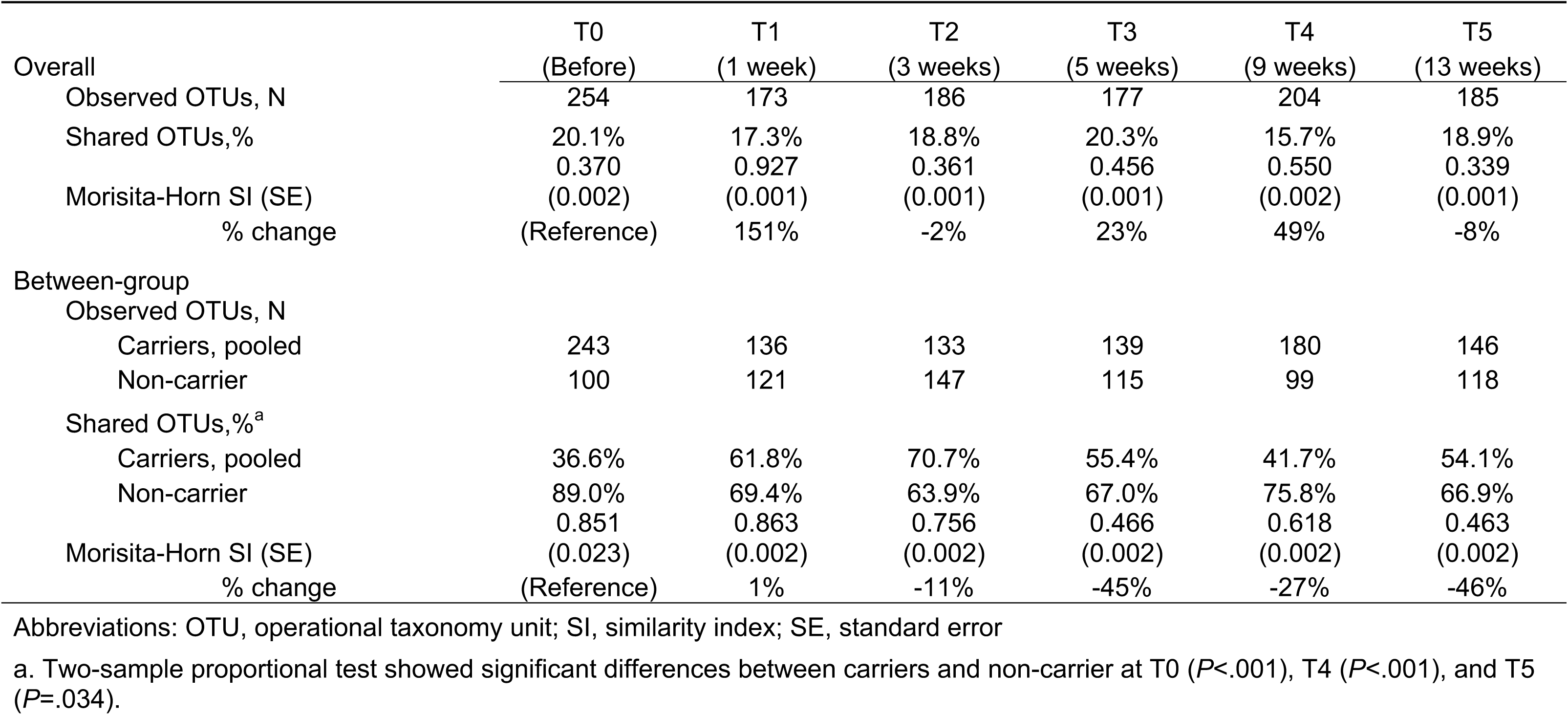
Community similarity estimates based on shared membership and community overlap by study visit and bewteen pretreatment carriers and the non-carrier

|  | T0 | T1 | T2 | T3 | T4 | T5 |
| --- | --- | --- | --- | --- | --- | --- |
|  | (Before) | (1 week) | (3 weeks) | (5 weeks) | (9 weeks) | (13 weeks) |
| Overall |  |  |  |  |  |  |
| Observed OTUs, N | 254 | 173 | 186 | 177 | 204 | 185 |
| Shared OTUs, % | 20.1% | 17.3% | 18.8% | 20.3% | 15.7% | 18.9% |
|  | 0.370 | 0.927 | 0.361 | 0.456 | 0.550 | 0.339 |
| Morisita-Horn SI (SE) | (0.002) | (0.001) | (0.001) | (0.001) | (0.002) | (0.001) |
| % change | (Reference) | 151% | -2% | 23% | 49% | -8% |
| Between-group |  |  |  |  |  |  |
| Observed OTUs, N |  |  |  |  |  |  |
| Carriers, pooled | 243 | 136 | 133 | 139 | 180 | 146 |
| Non-carrier | 100 | 121 | 147 | 115 | 99 | 118 |
| Shared OTUs, %<sup>a</sup> |  |  |  |  |  |  |
| Carriers, pooled | 36.6% | 61.8% | 70.7% | 55.4% | 41.7% | 54.1% |
| Non-carrier | 89.0% | 69.4% | 63.9% | 67.0% | 75.8% | 66.9% |
|  | 0.851 | 0.863 | 0.756 | 0.466 | 0.618 | 0.463 |
| Morisita-Horn SI (SE) | (0.023) | (0.002) | (0.002) | (0.002) | (0.002) | (0.002) |
| % change | (Reference) | 1% | -11% | -45% | -27% | -46% |

Abbreviations: OTU, operational taxonomy unit; SI, similarity index; SE, standard error
a. Two-sample proportional test showed significant differences between carriers and non-carrier at T0 ( $P<.001$ ), T4 ( $P<.001$ ), and T5 ( $P=.034$ ).

### Community dissimilarity and host factors

Based on eigenvalue partitioning, the initial staphylococcal carriage status (14.1%), changes in HNP1 expression levels (9.7%), and sampling time (7.6%) were major host factors for the observed variations in sample OTUs (Figure 2). As compared to a null model, the abovementioned covariates significantly explained 41% of the differences in the microbial community structure (*P*: .014), though the significance attenuated when within-group correlation was accounted for (*P*: .047). Clustering results were similar when using NMDS on dissimilarity metrics (data not shown).

**Figure 2.**
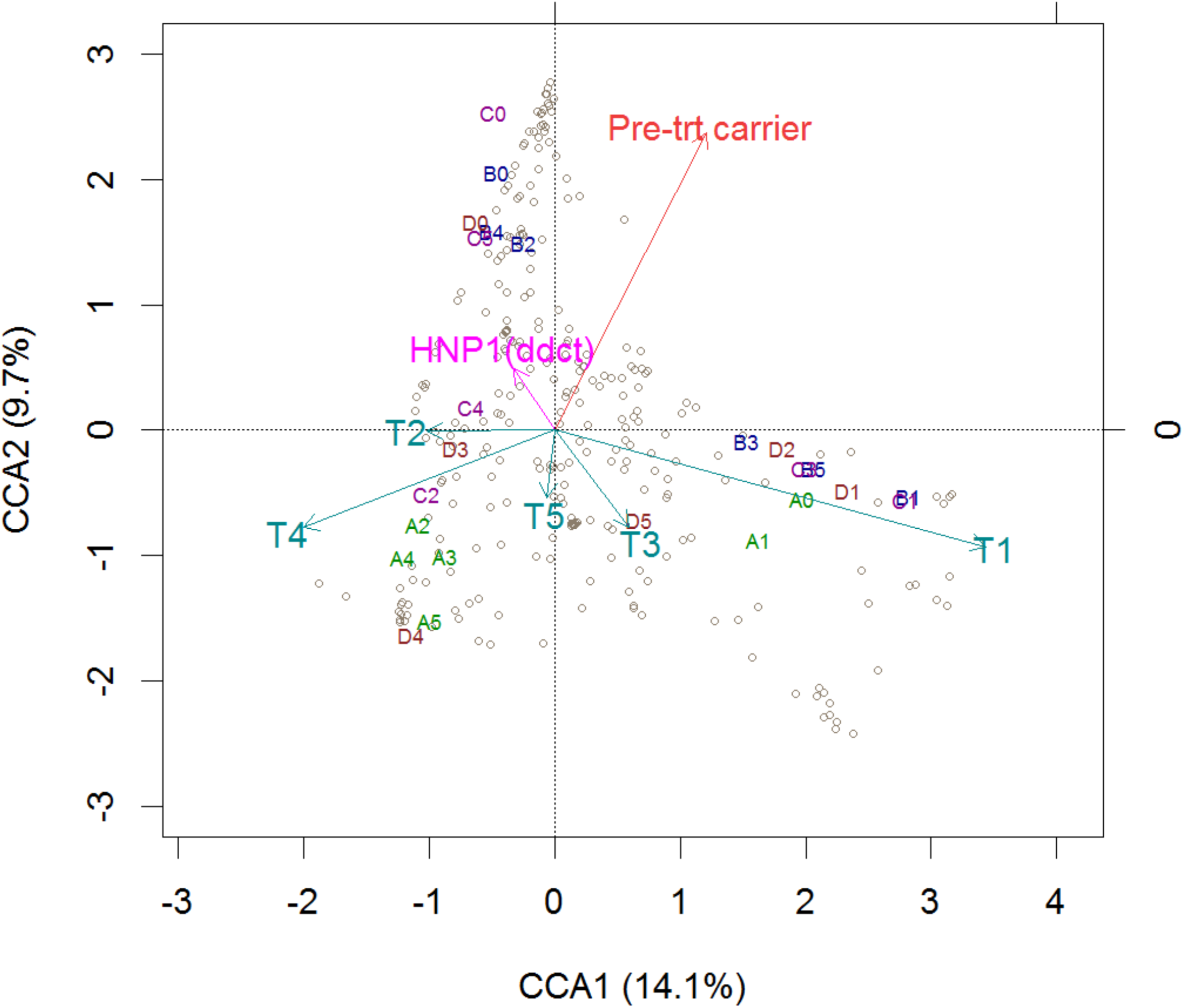
Results of multivariable-constrained correspondence analysis (CCA) showing the first two axes in grouping OTUs and samples by host factors, including the initial staphylococcal carriage status (“Pre-trt carrier”), levels of antimicrobial HNP1, and sampling times (Before, After, and 3 weeks, 5 weeks, 9 weeks, 13 weeks post-decolonisation). Letters denote individual study subjects and numbers specify sampling time points.

Although mupirocin use had a direct and specific depletion effect on *S. aureus* when comparing post-versus pre-decolonisation samples in differential abundance analysis (data not shown), levels of AMP expression were also determinants for sample dissimilarities (Supplementary Figure S3). When the baseline carriage status and levels of HNP1 expression were considered, besides Firmicutes, several genera of Actinobacteria and Proteobacteria were relatively depleted while a few Firmicutes and Actinobacteria were enriched in post-decolonisation samples as compared with those collected before mupirocin use (Figure 3).

**Figure 3.**
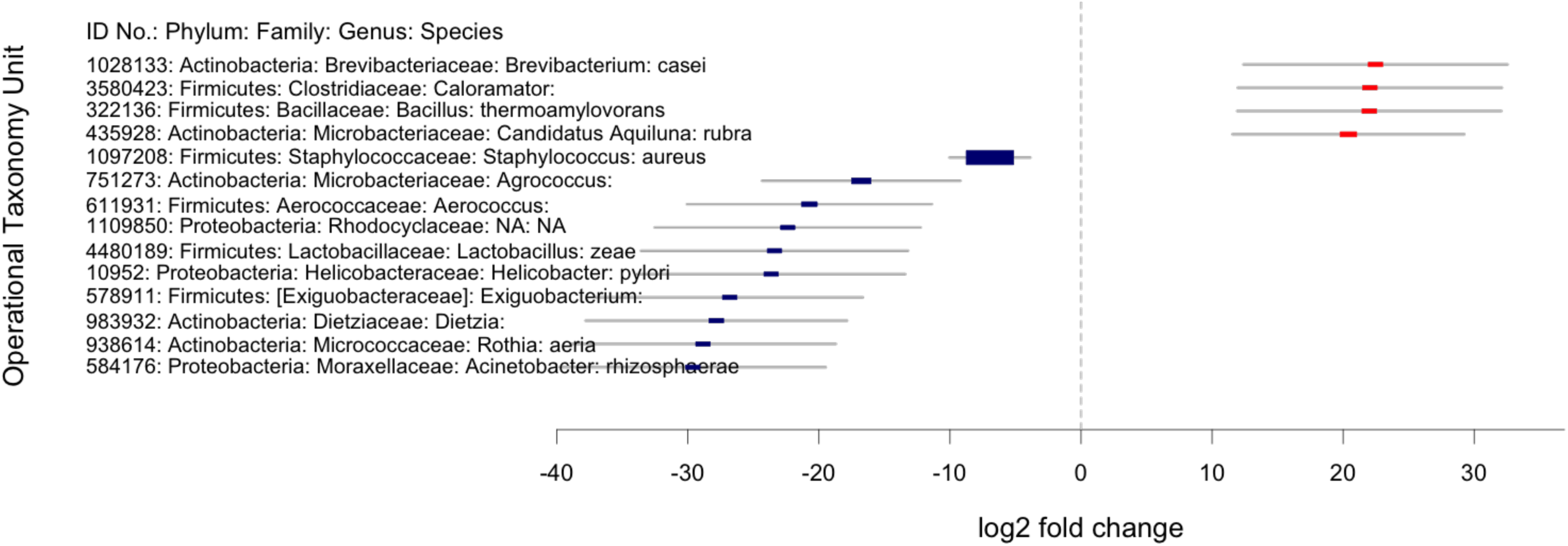
Results of differential abundance analysis showing specific OTUs that were substantially enriched (red) or depleted (blue) in samples collected after intranasal mupirocin use versus those collected before, with adjustment of expression levels of HNP1 and the baseline staphylococcal carriage status. (NA=unclassified)

## DISCUSSION

Despite of a small sample size, we took advantage of the serial sampling scheme in characterizing the temporal dynamics in the nasal microbiome after mupirocin decolonisation. Mupirocin initially reduced the microbial diversity up to 60%-75% of the baseline level before a noticeable resurgence of the community richness by 5 weeks post-decolonisation (T2). Although mupirocin has a well-recognized antimicrobial activity against staphylococci and streptococci,[30] its effects against the nasal commensals as a whole have not been formerly characterised. By examining the nasal microbiota longitudinally, mupirocin, along with the host antimicrobial responses, appeared to suppress a broad range of commensals, including several members in the phylum of Gram-negative Proteobacteria, effectively disrupting the ecological balance of the nasal microbiota both quantitatively and qualitatively. Although bacteria that appeared resistant to effects of the intranasal mupirocin and the host HNP1 were deemed mostly environmental in origin,[31-34] accumulating case reports suggested that *Brevibacterium casei*, once considered non-pathogenic, could lead to overt infections in high-risk individuals.[35-38] More research is needed to weigh the benefits of decolonisation against potential risks associated with the loss of colonisation resistance as observed in the gut microbiota.[39]

How mupirocin changed the microbial landscape generally depended on the initial staphylococcal carriage status; this finding was consistent with previous studies on how oral antibiotics could perturb the human gut microbiome.[40] Raymond and colleagues followed a cohort of 18 participants and six control subjects after a seven-day course of oral cefprozil (500mg) twice per day.[41] By comparing stool samples collected before (day 0) and after decolonisation (day 7 and day 90), the investigators concluded that the administered antibiotic had a predictable effect on all participants’ gut microbiota by their faecal microbiome on day 0.[41]

Investigators have shown that phenol-soluble modulins secreted by murine *Staphylococcus epidermis* (*S. epidermis*) could suppress the invasion of *S. aureus* and reduced the survival of Group A Streptococcus on mouse skin.[42] Intriguingly, we noted that higher levels of HBD3 and HNP1 were associated with the depletion and the enrichment of *C. kroppenstedtii*, respectively (Supplementary Figure S3), suggesting a potential role of the host immunity in determining the co-existent or mutually exclusive partnership among community members. We also found temporal variations in HBD3 levels, but not in HNP1, after mupirocin use; these findings were in line with the literature that HNP1 is constitutively produced by skin-resident neutrophils while the expression of HBD3 responds to exogenous stimuli to keratinocytes (such as bacterial colonisation).[16, 19, 43] Moreover, the observation that HNP1 levels were substantially higher in the (only) noncarrier than those in the carriers might be explained by a defective or down-regulated recruitment of neutrophils by staphylococcal carriers, possibly an immuno-modulating effect resulting from the host-pathogen interaction.[44] Although an early study by Cole et al. has suggested the otherwise,[45] we could infer that the abundance and the lack of *S. aureus* was possibly correlated with the local antimicrobial levels of HBD3 and HNP1 based on the observed associations between *C. kroppenstedtii* and expressed AMP levels.

### Limitations

The current pilot study was not well-powered and was mostly descriptive in nature. The fact that all study subjects were healthcare workers who routinely observed universal precautions at work further reduced the generalizability of our findings to healthy carriers in other settings. However, with repeated measures from the same individual, we have gained efficiencies while making intra-person (or intra-group) comparisons.

While the study findings suggested a distinction between sample microbiota from participants with or without *S. aureus* before decolonisation, we cautioned the interpretation of statistical comparisons by the initial carriage status as estimates associated with non-carriage were obviously under-estimated. Regardless, the current study contributed to the field by providing empirical estimates for future sample size and effect size calculation. Formal evaluations are required to better characterize mupirocin effects on the human nasal microbiome in absolutely noncarriers, intermittent as well as (strictly) persistent carriers.[12]

## CONCLUSIONS

In a small group of healthy staphylococcal carriers, we found short-term yet pervasive effects of mupirocin decolonisation on the ecology of the nasal microbiota. We also identified host immune correlates that were associated with substantial changes in the microbial community immediately after mupirocin use. These observations though preliminary warranted a larger-scale investigation to better understand how antibiotics may affect the second human genome and the host-pathogen interactions.

## List of abbreviations

AMP: antimicrobial peptide
CCA: constrained correspondence analysis
D_Shannon_: Shannon diversity index
D_Simpson_: Simpson diversity index
E_Shannon_: Shannon evenness index
E_Simpson_: Simpson evenness index
GAPDH: glyceraldehyde-3-phosphate dehydrogenase
HBD3: human beta-defensin 3
HNP1: human neutrophil peptide 1
IQR: interquartile range
MRSA: methicillin-resistant Staphylococcus aureus
NA: unclassified
NMDS: nonmetric multidimensional scaling
OTU: operational taxonomy unit
RNase7: ribonuclease 7
RT-qPCR: reverse transcription quantitative polymerase chain reaction
SE: standard error
SSI: surgical site infection
WK: week

## Ethics approval and consent to participate

The Institutional Review Board at Chang Gung Medical Foundation reviewed and approved the current study protocol and the participant consent form. Written informed consent was obtained from each study participant.

## Consent to publish

Not applicable.

## Availability of data and materials

Raw data and statistical codes that support results of the current study are available upon request.

## Authors’ contributions

SL conceived and designed the study, performed statistical analysis, and drafted the manuscript. YT helped manage the 16S rDNA sequence data and contributed to data analysis, results interpretation, and manuscript preparation. YL performed the laboratory work and assisted with the manuscript preparation. KC and CC participated in data interpretation and reviewed the manuscript. YH supervised the laboratory work, contributed to data interpretation, reviewed the manuscript. LYC involved in the study design, data analysis, and the manuscript drafting. All authors read and approved the final version of the manuscript.

## Acknowledgements

The authors are grateful to the study participants for their time support and commitment. The authors also wish to thank Ms. Ruby Lu for her laboratory and administrative support.

